# Vitamin D attenuates TNF-α-mediated neurotoxicity and improves functional recovery in experimental intracerebral haemorrhage

**DOI:** 10.1101/2025.06.22.660985

**Authors:** Stephen Yin Cheng, Karrie M. Kiang, Ziyi Zhong, Gilberto Ka-Kit Leung

**Author notes:** **Correspondence:** Professor Gilberto K.K. Leung, Department of Surgery, School of Clinical Medicine, LKS Faculty of Medicine, The University of Hong Kong, 21 Sassoon Road, Pokfulam, Hong Kong.

## Abstract

While the neuroprotective effects of vitamin D (Vit-D) have been demonstrated pre-clinically in a wide range of neurologic conditions, its potential use in the treatment of spontaneous intracerebral hemorrhage (ICH) has not been fully explored. We previously reported that Vit-D could expedite hematoma clearance in experimental ICH by inducing the conversion of M1 to M2 macrophage to enhance erythrophagocytosis^1,2^. Here, we provide new evidence on the dose-dependent effects of Vit-D on neuronal survival and functional recovery, lending further support for the clinical testing of Vit-D in the management of ICH.

## Results and Discussion

Adult male C57BL/6N mice with collagenase-induced ICH received Vit-D at 1000 IU/kg/day (low-dose) or 2000 IU/kg/day (high-dose) for 4 weeks, starting from 2-hour post-ICH. When compared with vehicle control, Vit-D treatment significantly improved motor function recovery on three independent assays, with the high-dose group outperforming the low-dose group **(Figure 1A)**. ICH upregulated poly (ADP-ribose) polymerase (PARP) and caspase-3 expression levels and caused extensive neuronal apoptosis in the peri-hematoma brain region **(Figure 1B)**, all of which were attenuated by Vit-D, resulting in fewer degenerated neurons compared to control.

**Figure 1.**
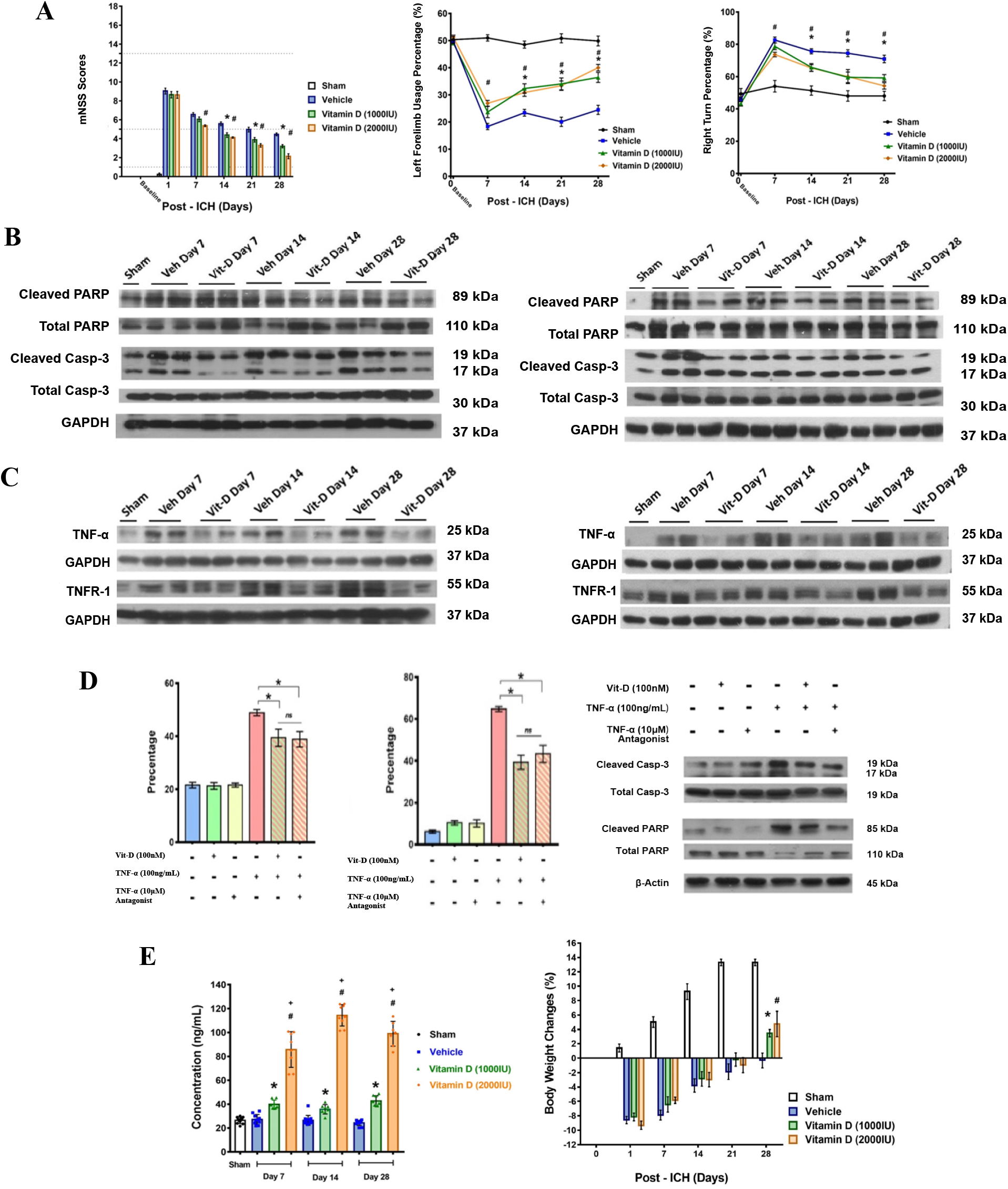
Effects of Vit-D supplementation on neurological and physiological outcomes in mice following ICH. **A.** Vit-D treatment (1000IU/day or 2000IU/day) was given to mice 2 hours post-ICH, and everyday post-ICH. Modified neurological function severity score (mNSS) assessment of mice at before and day 1, 7, 14, 21 and 28 post-ICH (left). Cylindrical test results of left forelimb usage: (left forelimb usage count + 1/2 both limbs usage) / (total forelimb usage count) × 100% (center). Corner turn test results of percentage right turn: (right turn count) / (total turn count) × 100% (right). Error bars indicate mean ± SEM. Two-way ANOVA followed by Tukey’s multiple comparison test. **B**. Western blot measurement of Caspase-3 and PARP protein expression after ICH. Vit-D 1000IU/kg (left) and 2000IU/kg (right) was given to post-ICH mice 2 hours post-ICH, and everyday post-ICH. **C**. Western blot measurement of TNF-α and TNFR1 protein expression before ICH and at day 7, 14 and 28 post-ICH. The expression of GAPDH is used as loading control. Vit-D 1000IU/kg (left), Vit-D 2000IU/kg (right) or drug vehicle was given to post-ICH mice daily by oral gavage. **D**. TNFR1 protein expression in neurons measured by flow cytometry analysis (left). Apoptotic cell population in neurons measured by Annexin-V-PI flow cytometry analysis (center). Western blot measurement of Caspase-3 and PARP protein expression. Cell and protein were harvested from the cultured mouse embryonic cortical neurons 48 hours after exposure to Vit-D (100nM), TNF-α (100ng/mL), TNF-α antagonist (10μM). **E**. Serum 25(OH)D level measured by ELISA analysis on blood harvested from mice before ICH, and at day 7, 14, 28 post-ICH (left). ^*^Vit-D 1000IU/kg vs Vehicle; ^#^Vit-D 2000IU/kg vs Vehicle; ^+^Vit-D 1000IU/kg vs Vit-D 2000IU/kg. Mice body weight changes were recorded in different experimental groups from mice before ICH (day 0) and at day1, 7, 14 and 28 post-ICH (right). Error bars indicate mean ± SEM. One-way measures ANOVA and Tukey’s multiple comparison test.

Tumor necrosis factor-alpha (TNF-α) is a proinflammatory cytokine responsible for ICH-induced secondary brain injury. The release of toxic hemoglobin metabolites in ICH triggers the production of TNF-α by microglia and astrocytes, causing additional neurovascular injury and poorer patient outcome.^3^ We found that Vit-D treatment could downregulate the expression of TNF-α *in vivo*, achieving almost complete inhibition at high-dose within the affected cerebral hemisphere (**Figure 1C**). We therefore examined the neuronal expression of TNF receptor-1 (TNFR1), which was found to reach its peak by day 14 post-ICH and was significantly downregulated by Vit-D treatment (**Figure 1C**). Mechanistic studies using primary culture of embryonic cortical neurons showed that *in vitro* treatment with exogenous TNF-α increased the neuronal expression of TNFR1 from 20% at baseline to 50%, and the percentage of apoptotic neurons from 6-10% at baseline to over 60% **(Figure 1D)**. The addition of Vit-D or a TNF-α antagonist significantly reduced the expression levels of both TNF-α and TNFR1, together with the downregulation of caspase-3 and PARP expression levels and a reduction in neuronal apoptosis (**Figure 1D**). The findings indicate the ability of Vit-D to act at both the ligand and the receptor expression levels of TNF-α-mediated cell death, potentially giving rise to more robust neuronal protection.

We also examined the adverse effects of hypervitaminosis D. When compared with control, low-dose and high-dose Vit-D treatments increased serum Vit-D level by 150% and 350%, respectively, with the high-dose treatment yielding a serum level of 120 ng/ml (**Figure 1E**). No changes in serum calcium level occurred, and Vit-D-treated animals were able to regain body weight at a significantly higher rate when compared with control (**Figure 1E**). In human adults, Vit-D toxicity is thought to occur at a serum Vit-D level of 150 ng/ml, commonly as the result of months of excessive intake; our high-dose regimen, administered for a relatively short duration, is therefore likely to be well-tolerated ^4^. However, individuals with Vit-D hypersensitivity or excessive calcium intake may react differently; close monitoring of Vit-D and calcium serum levels is therefore necessary. The situation is further complicated by the high prevalence of Vit-D deficiency in certain populations. Another study of our group showed that Vit-D deficiency was associated with poorer outcomes in mice post-ICH, while a single dose of Vit-D could provide significant improvement, suggesting that individuals with pre-existing Vit-D deficiency may require an even high dosage of Vit-D treatment in order to obtain the maximal therapeutic benefit ^5^. Further clinical studies are needed to ascertain the optimal, safe dosage for each individual patient.

In conclusion, pre-clinical studies have shown that Vit-D supplementation could significantly improve the outcome of ICH, at least partially through the suppression of TNF-α-mediated neuronal cell death. Clinical translation of these findings should adopt a personalized approach based on point-of-care measurement of serum calcium and Vit-D levels to cater for pre-existing Vit-D status and varied responses to Vit-D treatment.

## References

1. Liu, J., Zhu, Z. & Leung, G. K.-K. Erythrophagocytosis by Microglia/Macrophage in Intracerebral Hemorrhage: From Mechanisms to Translation. Front. Cell. Neurosci. 16, 818602 (2022).

2. Liu, J. et al. Vitamin D Enhances Hematoma Clearance and Neurologic Recovery in Intracerebral Hemorrhage. Stroke 53, 2058–2068 (2022).

3. Hua, Y. et al. Tumor Necrosis Factor-α Increases in the Brain after Intracerebral Hemorrhage and Thrombin Stimulation: Neurosurgery 58, 542–550 (2006).

4. Marcinowska-Suchowierska, E., Kupisz-Urbańska, M., Łukaszkiewicz, J., Płudowski, P. & Jones, G. Vitamin D Toxicity–A Clinical Perspective. Front. Endocrinol. 9, 550 (2018).

5. Chan, A. A. et al. Acute calcitriol treatment mitigates vitamin D deficiency-associated mortality after intracerebral haemorrhage. Neurosci. Lett. 838, 137922 (2024).

